# Transcriptome analysis of *Culter alburnus* gonad tissues for discovery of sex-related genes

**DOI:** 10.1101/351759

**Authors:** Jianbo Zheng, Yongyi Jia, Shili Liu, Wenping Jiang, Meili Chi, Shun Cheng, Zhimin Gu

## Abstract

*Culter alburnus* is an important commercially fish species for freshwater breeding in China, and the females grow faster than the males. However, the molecular genetic mechanism of sex determination in *C.alburnus* is still poorly characterized. Here, we performed *de novo* assembly of a transcriptome from adult fish tissues of different gender using short read sequencing technology (Illumina). Our results showed that a total of 364,650 unigenes using Trinity software were obtained, giving rise to an average of 561.92 bp per read. Among them, 70,215 sequences matched known genes, including 5,892 male-biased unigenes and 942 female-biased unigenes. Many sex-related genes and pathways were identified based on annotation information. These results would provide new insights into the genetic mechanism of *C.alburnus* sex determination and also establish an important foundation for further research on aquaculture breeding.

---

*Culter alburnus* is one kind of important fish species for freshwater breeding in China, and the female individual grows faster than the male one. Thus, developing sex control technology to produce all female population can improve breeding yield and increase economic income. Although we succeeded to obtain all female species through gynogenesis and sex reversal, the genetic mechanism of sex determination in *C.alburnus* is still poorly characterized. These remaining problems severely limit sex control breeding technology application in *C.alburnus*.

Recently, with advancements in next-generation sequencing technologies, transcriptome profiling provides an invaluable tool for gene discovery and functional analysis [1]. So far, several fish species that involved in sex determination and differentiation by RNA-seq are reported, including *Oreochromis niloticus* [2], *Xiphophorus maculatus* [3], *Ictalurus punctatus* [4], *Paralichthys olivaceus* [5], *Danio rerio* [6], *Oncorhynchus mykiss* [7], and so on. These transcriptome sequencing data with identification of the sex-differentially expressed genes is particularly helpful to understand and elucidate the gene regulatory mechanism of sex determination and differentiation.

Previous studies on *C.alburnus* have mainly focused on embryonic development, growth and reproduction, little involved in the levels of molecular genetic mechanism. In this study, we performed *de novo* assembly of a transcriptome from adult fish tissues of different gender using short read sequencing technology (Illumina). These assembled, annotated transcriptome sequences and gene expression profiles would offer a rapid approach to identify key candidate regulatory genes underlying the processes of gonad development. Moreover, demonstrating of the regulatory mechanisms associated with sex determination will be essential for accelerating and advancing aquaculture breeding programs on *C.alburnus* in the future.

## Materials and Methods

### Fish samples preparation and RNA extraction

The fish (three males and three females) used in this study were provided from the Balidian breeding base of Zhejiang Institute of Freshwater Fisheries (Huzhou, Zhejiang Province). The tissues of testes and ovaries were immediately frozen in liquid nitrogen, and stored at −80°. Total RNA was extracted by SV Total RNA Isolation System (Promega) and was treated with DnaseI (Ambion, USA) to remove contaminative DNA.

### Library construction and Illumina sequencing

cDNA library was generated by the SMART cDNA library construction kit (Clontech, USA) according to the manufacturer’s protocol. Firstly, mRNA was sheared with fragmentation buffer to produce short bands (200 bp), from which the first-strand DNA was synthesized using random primers and reverse transcriptase. Subsequently, the synthesized cDNA was subjected to end-repair by adding End Repair Mix, ‘A’ base addition and ligated to adapters according to Illumina’s library construction protocol. Finally, the completed libraries were sequenced on the Illumina HiSeq 2000 platform [8, 9].

### Sequence data analysis and assembly

The raw reads were initially pre-processed by removing adaptor sequences, low quality sequences, empty reads, and higher N rate sequences to obtain the high quality clean data using SeqPrep (https://github.com/jstjohn/SeqPrep) and Sickle (https://github.com/najoshi/sickle). The trimmed and size-selected reads were then *de novo* assembled by Trinity program (http://trinityrnaseq.sourceforge.net/) [10].

### Gene annotation and function classification

All assembled unigenes were compared against the protein databases, such as non-redundant (NR), Kyoto Encyclopedia of Genes and Genomes (KEGG), Search Tool for the Retrieval of Interacting Genes (String), Swissprot and Pfam (Swissprot), to obtain function annotations using BlastX with a typical cut-off E-value<1e^−5^. GO annotation was performed using Blast2GO (http://www.blast2go.com/b2ghome), Clusters of Orthologous Groups (COG) (http://www.ncbi.nlm.nih.gov/COG/) andKEGG (http://www.genome.jp/kegg/) were used to predict possible functional classifications and metabolic pathways [9].

### Identifcation of sex-related differentially expressed genes

RPKM (Reads Per Kilobase of exon model per Million mapped reads) was directly used to compare the difference of gene expression level between male and female individuals [11]. The formula was as follows:

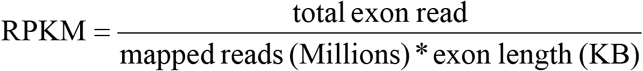

FDR<0.05 and log_2_|FC|≧1 were set as the calculation criterion for significantly differential expression analysis. Based on searching sex-related keywords and other published strategies were to predict sex differentially expressed genes.

### KEGG enrichment analysis of differentially expressed genes

KEGG (Kyoto Encyclopedia of Genes and Genomes) was a public database for revealing high level genomic function [12]. In this study, the KEGG functional enrichment analysis of differential expressed genes was tested by KOBAS software (http://kobas.cbi.pku.edu.cn/home.do) [13,14].

## Results

### Sequencing and *de novo* transcriptome assembly of *C.alburnus*

In order to identify sex-related genes and clarify the mechanism of sex determination in *C.alburnus*, two cDNA libraries were constructed from testes and ovaries for transcriptome analysis. After eliminating adapter sequences and filtering out the low-quality reads, the Illunima HiSeq sequencing produced 127,931,976 raw reads from the testes and 27,215,890 raw reads from the ovaries, respectively (Table 1). The *de novo* assembly of high quality transcriptomic reads generated 364,650 unigenes using Trinity software and the lengths were distributed as 561.92 bp, 22,423 bp, 224 bp, 640 bp, 266 bp from average, longest, shortest, N50 and N90 length,respectively (Fig 1 a). In addition, most of these unigenes (90.47%) were distributed in the 200-1000 bp region (Fig 1 b).

**Table 1.**
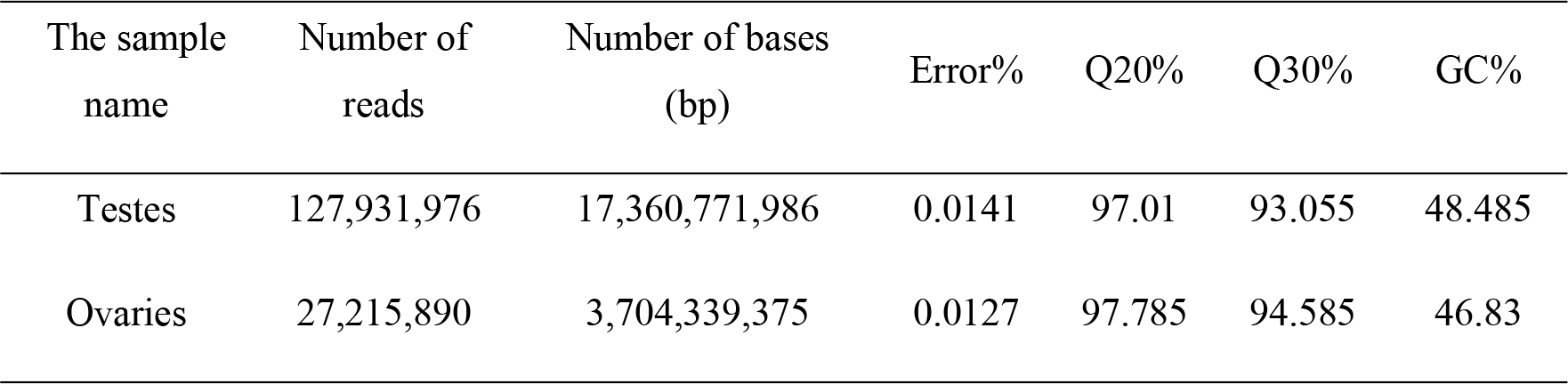
Statistics of the sequencing results from testes and ovaries of *Culter alburnus*.

**Fig 1.**
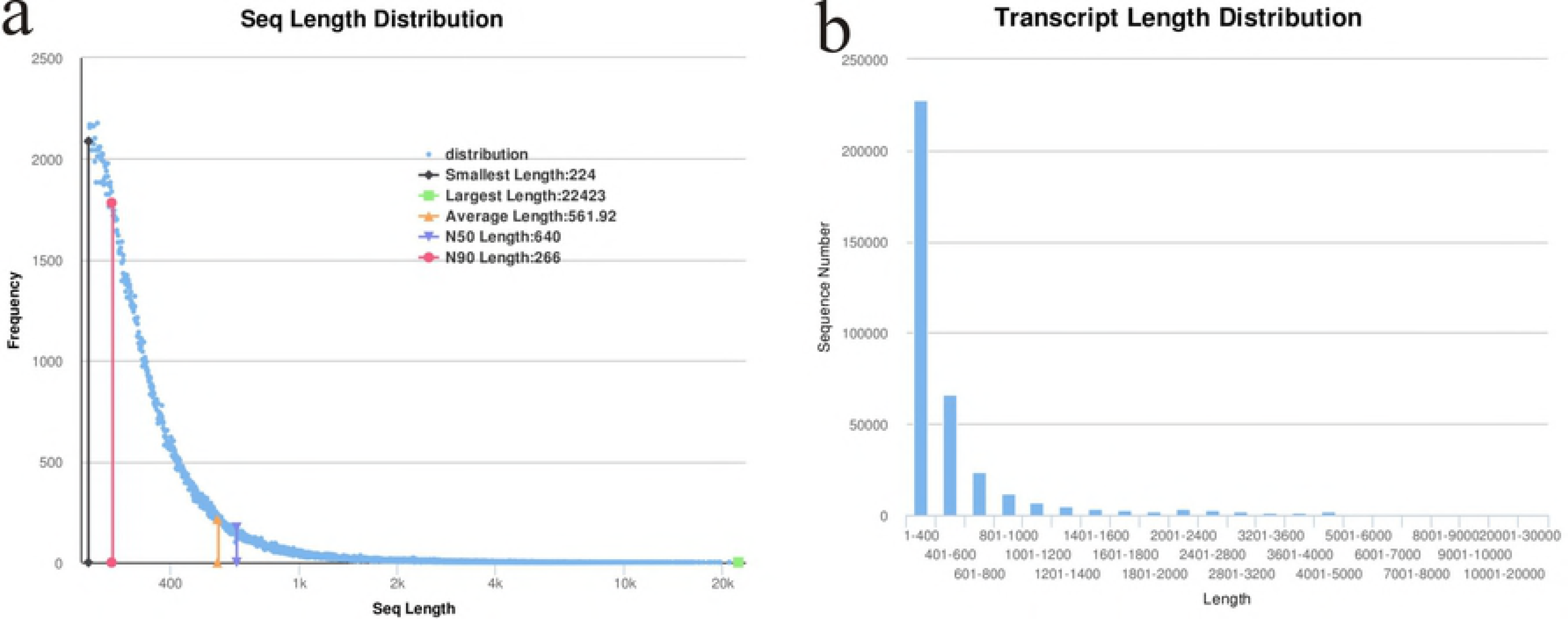
Length distribution of unigenes assembled by Trinity. (a) Sequence length distriution from average, longest, shortest, N50 and N90 length, respectively. (b) Size distribution for transcript sequence.

### Function annotation, classifictiaon and bioinformatical analysis

The unigenes were initially processed by searching the non-redundant protein databases using BlastX and a total of 70,215 sequences (19% of unigenes) matched known genes. Among those distinct sequence, the majority of sequences (30,379) had strong homology with *Danio rerio*, followed by *Oncorhynchus mykiss* (3,190), *Astyanax mexicanus* (2,481), *Mus musculus* (1,445), *Tetraodon nigroviridis* (784), *Stegastes partitus* (767), and other or unknown species, which made up 44.4% of total genes (Figure 2).

**Fig 2.**
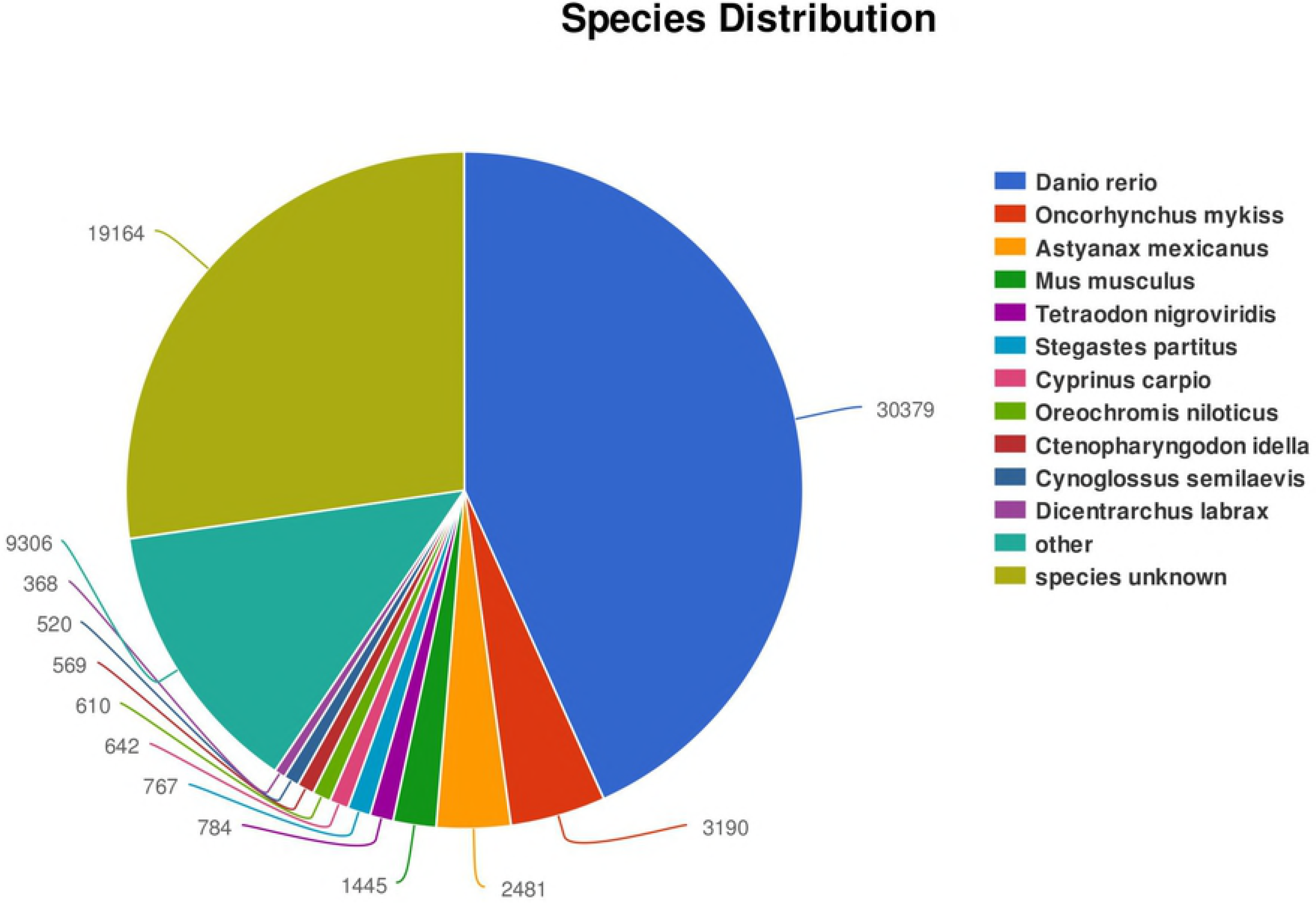
Species distribution that match to the sequences of *C.alburnus*. Each sector represents a species. The number of homologous sequence corresponding each species using BlastX are indicated near the sector.

Gene Ontology (GO) and Cluster of Orthologous Groups of proteins (COG) were used to predict and classify possible functions of the unigenes. For GO analysis, all annotated unigenes (30,654) were classified into three functional categories, including biological process, cellular component and molecular function (Fig 3). In biological process, genes were divided into 25 classifications and cellular process (62.34%) was the largest subcategory. In cellular component, genes involved in cell (44.5%) and cell part (44.5%) were the most abundant. In molecular function, genes were grouped into 20 classifications and the most represented molecular function were bingding (57.8%). For COG analysis, 20,349 unigenes were annotated and divided into 25 specific categories (Fig 4). The most common category was ‘general function prediction only’ with 3,974 unique sequences, followed by ‘Signal transduction mechanisms’ (2,619), ‘Posttranslational modification, protein turnover, chaperones’ (1,876), ‘Replication, recombination and repair’ (1,239). ‘Cell motility’ (15), ‘Extracellular structures’ (0), ‘Nuclear structure’ (13) were the smallest COG categories.

**Fig 3.**
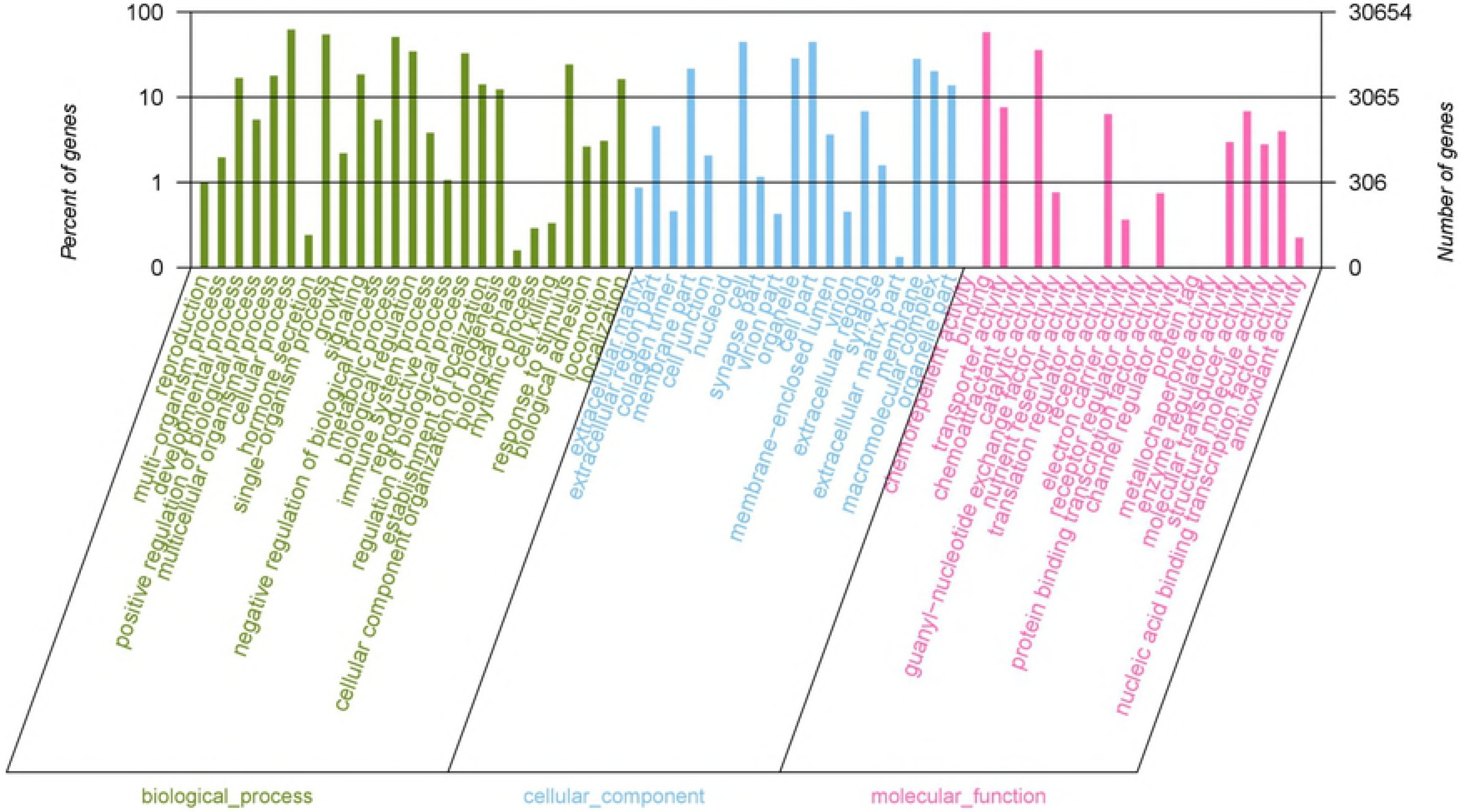
Gene ontology (GO) category for the transcriptome of *C.alburnus*. Green represents biological process, blue represents cellular component, pink represents molecular function.

**Fig 4.**
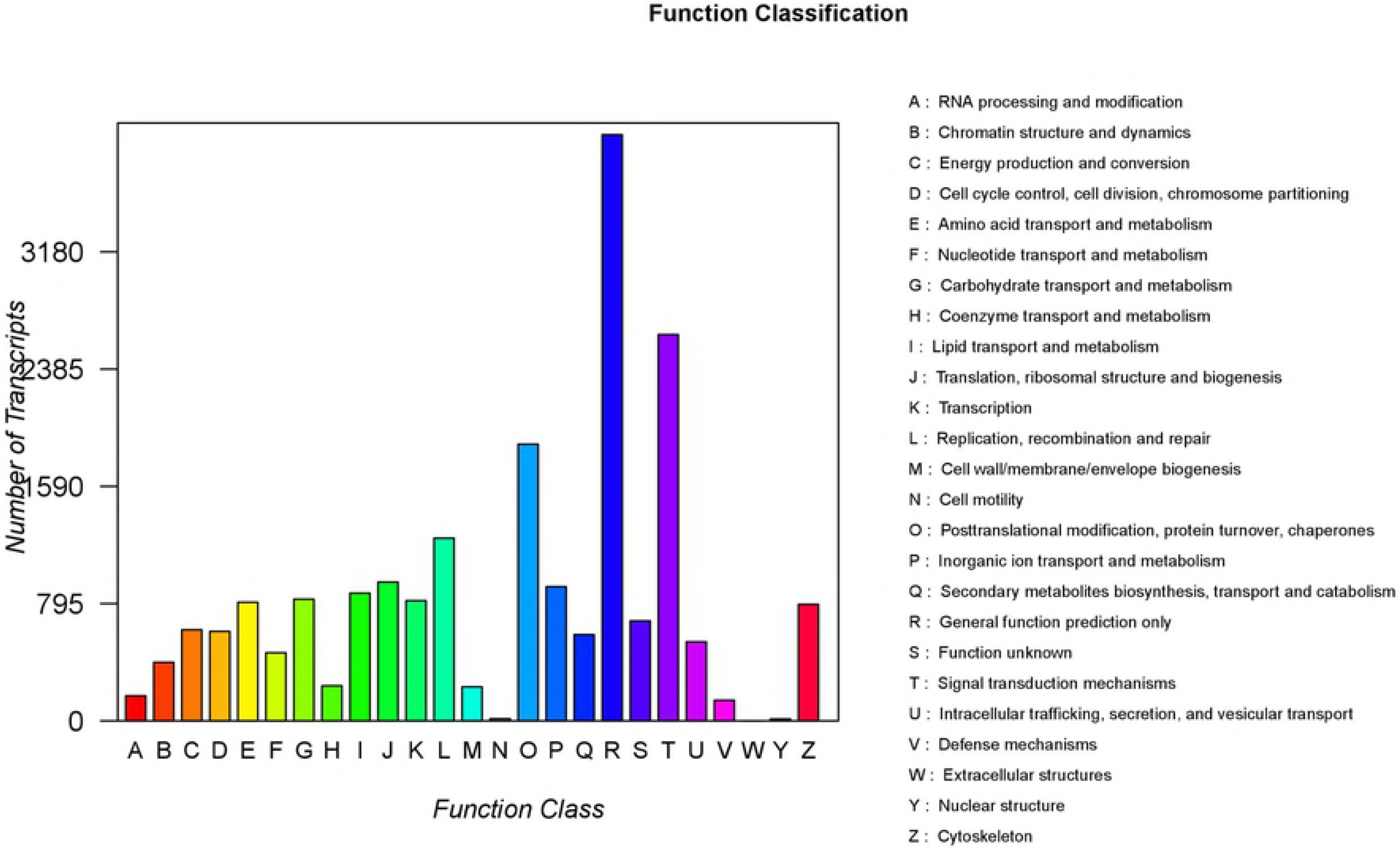
Clusters of orthologous groups (COG) Classification and annotation for *C.alburnus* unigenes. Each color column represents the functional classification of a COG, which indicated by the capital letter A-Z.

To systematically understand the biological functions of genes, the unigene metabolic pathway analysis was conducted using the Kyoto Encyclopedia of Genes and Genomes (KEGG) annotation system. According to the KEGG results, We mapped 29,467 unigenes to 362 KEGG pathways (Fig 5). ‘Pathways in cancer’ pathway had the largest number of unigenes (1,233), followed by ‘PI3K-Akt signaling pathway’ (1,022), ‘Calcium signaling pathway’ (921), ‘Neuroactive ligand-receptor interaction’ (921), ‘MAPK signaling pathway’ (912) and ‘cAMP signaling pathway’ (883).

**Fig 5.**
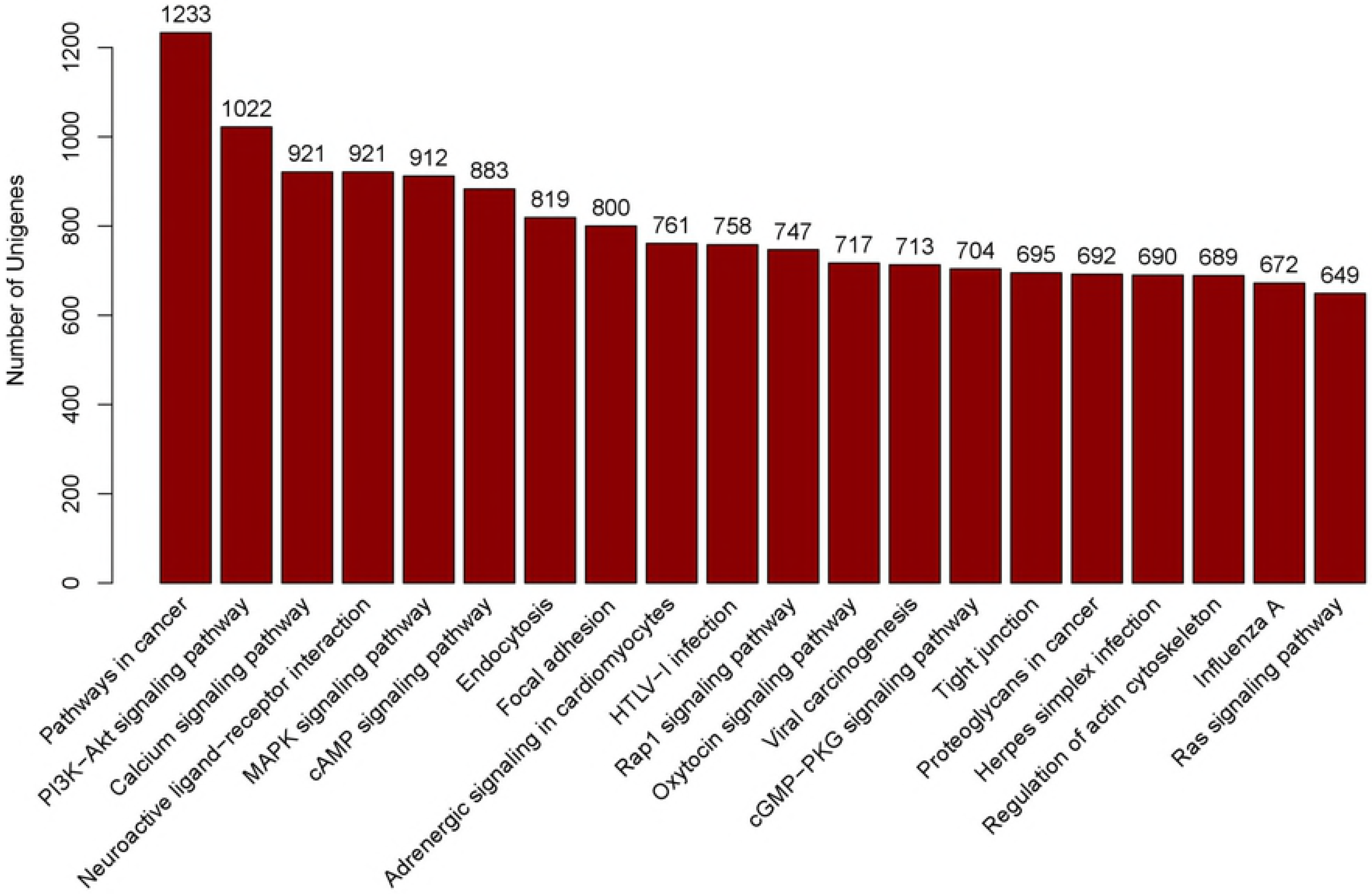
The top twenty pathways with most abundant unigenes.

### Sex-related genes idetification from comparison of gonad tissues transcriptomes

Gene differential expression analysis was performed by edgeR software, which generated gene read count for differential expression calculation [15]. Based on calculation criterion of FDR<0.05 and log2|FC|≧1, 38,167 unigenes showied signifcant expression difference between male and female individuals, including 5,892 (15.44%) male-biased unigenes and 942 (2.47%) female-biased unigenes (Fig 6). According to our sequence analysis and other published search strategies [5], sex-related well-documented genes were identifed (Table 2). Obviously, many sex-linked genes, especially those located upstream of gonadal development regulatory network, had much more significant difference trend in expression levels, such as *dmrt1*, *foxl2*, *cyp19a*, and so on.

**Fig 6.**
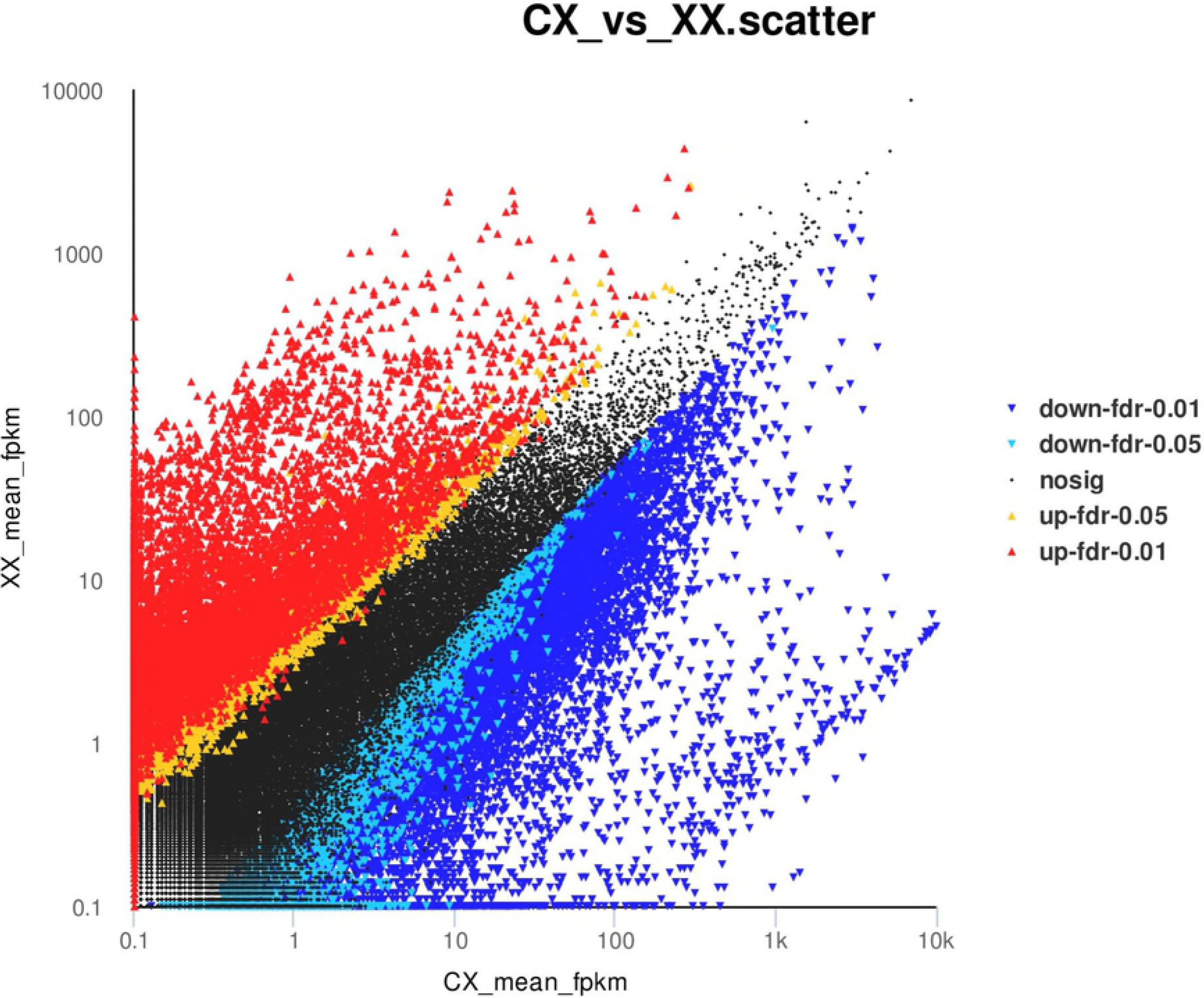
Comparisons of gene expression between females and males of *C.alburnus*. XX represents male individuals, CX represents female individuals.

**Table 2.**
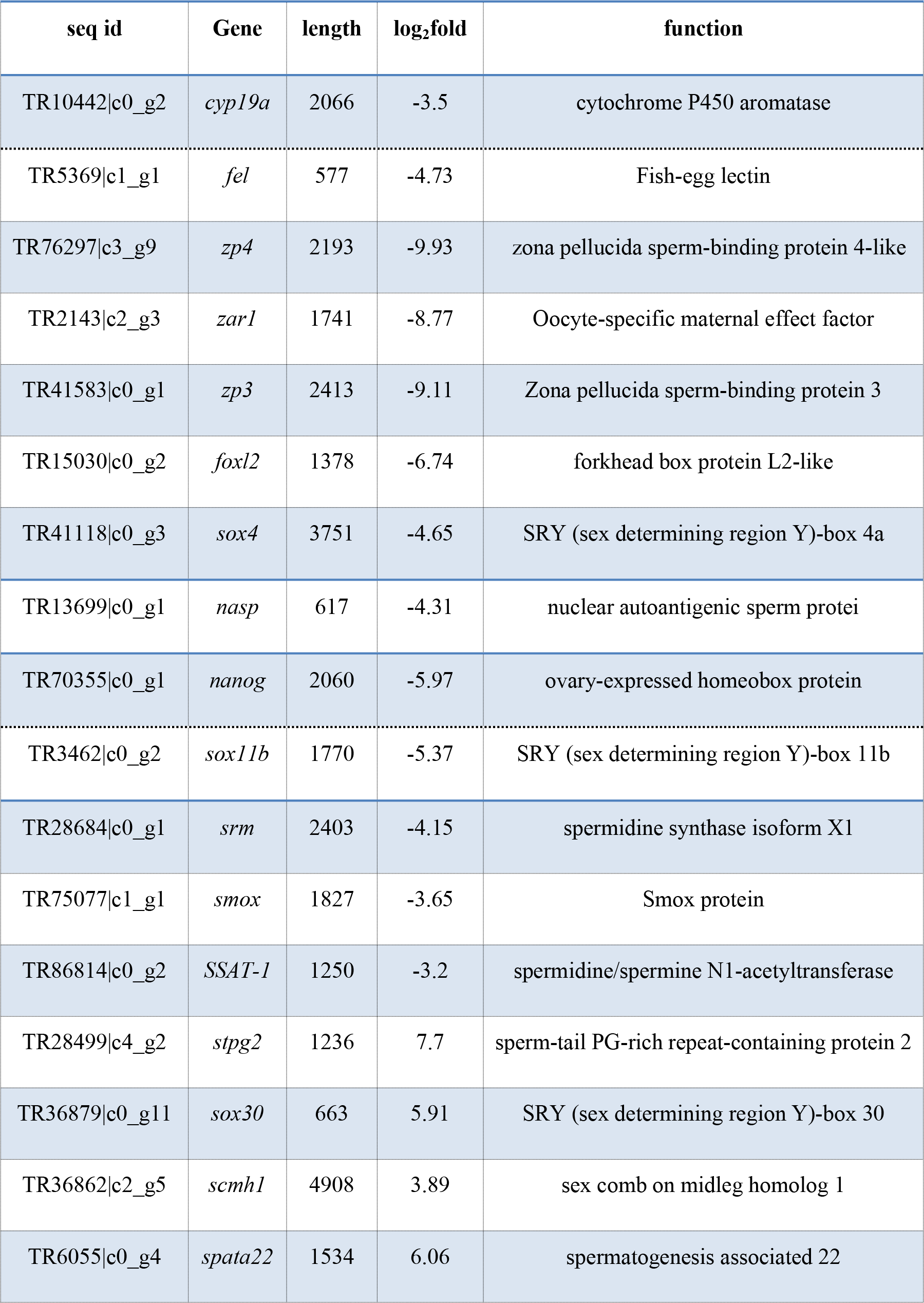
Representative sex-related differentially expressed genes in *C.alburnus*.

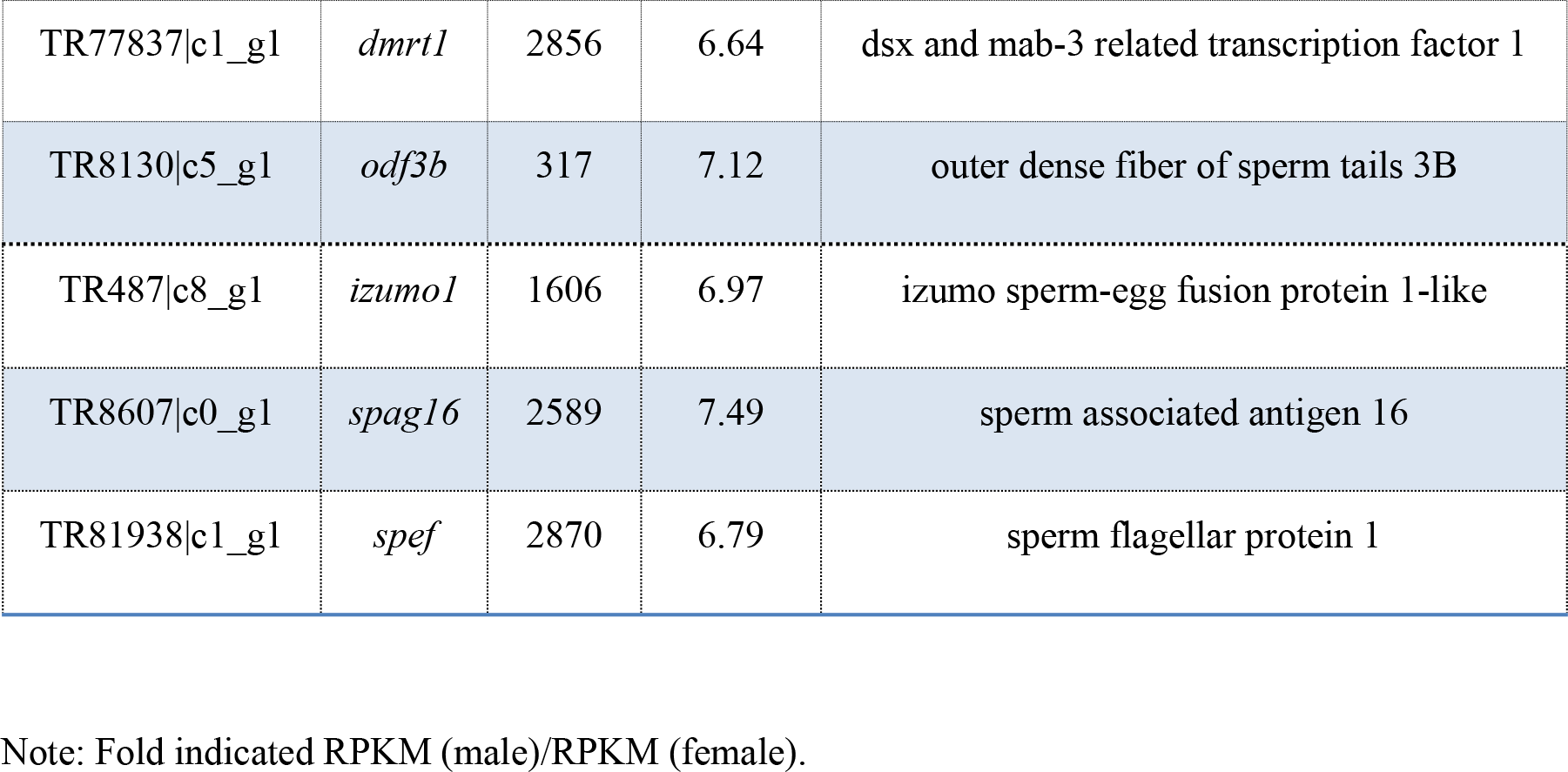

### KEGG enrichment analysis of sex-differential expression genes

KEGG enrichment analysis was applied in revealing high level genomic function to discover the metabolic processes and signal transduction pathways. Statistic and analysis of sex-differential expression genes involving in different category of metabolic pathways, found that these genes mainly enriched in Pathways in cancer, Calcium signaling pathway, cAMP signaling pathway, PI3K-Akt signaling pathway, Endocytosis, MAPK signaling pathway, Neuroactive ligand-receptor interaction, cGMP-PKG signaling pathway. Among them, several pathways associated with gonadal development and sex maintenance were identified, such as ovarian steroidogenesis, estrogen signaling pathway, progesterone-mediated oocyte maturation, prolactin signaling pathway, oocyte meiosis, TGF-beta signaling pathway, steroid hormone biosynthesis.

## Discussion

Previous study with identification sex-linked genetic markers using AFLP analysis (amplified fragment length polymorphism) in our lab showed that *C.alburnus* sex differentiation may be controlled by strict genetic regulation. However, the role of genetic factors in sex determination and differentiation is unclear and little evidence can demonstrate its genetic mechanism. Therefore, the discovery of sex-related genes and biological regulatory pathways would provide important clues for systematic elucidation sex determination mechanism of *C.alburnus*.

In this study, we performed the *de novo* transcriptome sequencing of *C.alburnus* testes and ovaries tissues by using Illumina Hiseg platform [8,16]. A total of 364,650 unigenes using Trinity software were obtained, giving rise to an average of 561.92 bp per read. About 19% of unigenes were found to match known genes using the public databases and the majority of sequences showed the greatest similarity to *Danio rerio*.

There has several reasons for explaining the rest of unigenes without annotations: i) most of these unigenes are short fragments; ii) these unigenes are probably new genes or unique to the species of *C.alburnus*. iii) these genes may be non-coding RNA sequences [17]. Hence, these transcriptome data would provide a useful resource for future genetic or genomic studies on this species.

It is known that the mainly aim of transcriptome sequencing is to identify a large number of candidate genes potentially involved in specific biological processes, such as growth, reproduction, and gonad development. In present study, our results revealed large quantities of sex-biased genes that showed sexual dimorphism between ovary and testis by analysis the gene expression profiles of *C.alburnus* gonads. Moreover, multiple sex-related genes were identifed to be involved in several biological pathways associated with gonadal development and sex maintenance, including ovarian steroidogenesis, estrogen signaling pathway, Gn RH signaling pathway, oocyte meiosis, TGF-beta signaling pathway, steroid hormone biosynthesis,and Wnt signaling pathway.

Taken together, this was the first attempt to perform RNA-seq technology to identify differentially expressed genes between ovaries and testes on *C.alburnus*. Additionly, a significant number of sex-related biological pathways associated with the unique sequences were found. These results would provide new insights into the genetic mechanism of *C.alburnus* sex determination and also establish an important foundation for further research on aquaculture breeding.

## Conflicts of interest

The authors declare no conflicts of interest.

## Acknowledgements

This work was financially supported by grants from, Natural science foundation for young scientists of Zhejiang province (LQ18C190001), Zhejiang Science and Technology Major Program (2016C02055-1), and Natural Science foundation of Huzhou (2017YZ02).

